# Design and evaluation of a large sequence-capture probe set and associated SNPs for diploid and haploid samples of Norway spruce (*Picea abies*)

**DOI:** 10.1101/291716

**Authors:** Amaryllis Vidalis, Douglas G. Scofield, Leandro G. Neves, Carolina Bernhardsson, María Rosario García-Gil, Pär K. Ingvarsson

## Abstract

Massively parallel sequencing has revolutionized the field of genetics by providing comparatively high-resolution insights into whole genomes for large number of species so far. However, whole-genome resequencing of many conspecific individuals remains cost-prohibitive for most species. This is especially true for species with very large genomes with extensive genomic redundancy, such as the genomes of coniferous trees. The genome assembly for the conifer Norway spruce (*Picea abies*) was the first published draft genome assembly for any gymnosperm. Our goal was to develop a dense set of genome-wide SNP markers for Norway spruce to be used for assembly improvement and population studies. From 80,000 initial probe candidates, we developed two partially-overlapping sets of sequence capture probes: one developed against 56 haploid megagametophytes, to aid assembly improvement; and the other developed against 6 diploid needle samples, to aid population studies. We focused probe development within genes, as delineated via the annotation of ~67,000 gene models accompanying *P. abies* assembly version 1.0. The 31,277 probes developed against megagametophytes covered 19,268 gene models (mean 1.62 probes/model). The 40,018 probes developed against diploid tissue covered 26,219 gene modules (mean 1.53 probes/model). Analysis of read coverage and variant quality around probe sites showed that initial alignment of captured reads should be done against the whole genome sequence, rather than a subset of probe-containing scaffolds, to overcome occasional capture of sequences outside of designed regions. All three probe sets, anchored to the *P. abies* 1.0 genome assembly and annotation, are available for download.

## Introduction

Massively parallel sequencing has revolutionized the field of genetics by providing comparatively high-resolution insights into whole genomes for large number of species so far. However, whole-genome resequencing of many conspecific individuals for the assessment of genetic variation in large-scale population studies (e.g., Wang et al. 2016), or for linkage-based studies such as association mapping or genomic breeding (e.g., Wang et al. 2017) remains cost-prohibitive for most species. This is especially true for species with very large genomes with extensive genomic redundancy, such as the genomes of coniferous trees (Nystedt et al. 2013, Neale et al. 2014). A number of methods have recently been developed to overcome this problem, focusing on reducing genome complexity to allow partial sequencing of whole genomes. The genomic regions sequenced by these methods are either anonymous, if based on reduced representation libraries generated by restriction enzymes (e.g., Davey and Blaxter 2010), or targeted, using primers and/or probes targetting selected genomic regions for high-throughput amplification or capture for later sequencing (e.g., Clark et al. 2011).

Sequence capture is a targeted reduced-representation method that can maximize the advantage of additional available genomic information such as a reference genome and associated annotation to target, extract and sequence selected regions of a genome, usually with the aim to conduct comparative analysis across several individuals. Sequence capture is a hybridization-based technique which shears genomic DNA and uses synthetic oligonucleotide probes to hybridize with fragments corresponding to specific regions within the genome, which are then captured and sequenced for further analysis. Depending on the hybridization technology, varying numbers of probes can be used. For humans, multiple technologies are available which contain probes sufficient to capture whole exomes (Clark et al., 2011; Shigemizu et al., 2015). Such comprehensive approaches can be used successfully in model species with well-annotated genomes (Fu et al., 2013; Zhou et al. 2012; Zhou et al. 2014). However, because sequence capture relies largely on the accuracy of genome annotations and the uniqueness of probe targets, it may exhibit reduced efficiency when applied to non-model species with incomplete annotations and/or species with complex genomes containing much repetitive content (Neves et al., 2013; Suren et al., 2016). Thus, we chose a sequence capture technology that had been used successfully in large, repeat-rich, relatively uncharacterized plant genomes (Rapid Genomics Capture-Seq; Neves et al., 2013).

The genome assembly for the conifer Norway spruce (*Picea abies*) was the first published draft genome assembly for any gymnosperm (Nystedt et al. 2013). From a total genome size estimated to be 19.6 Gbp, the *P. abies* genome version 1.0 included 12 Gbp in scaffolds larger than 200 bp with 4.3 Gbp in scaffolds larger than 10 kbp. Our overall goal was to develop a dense set of genome-wide SNP markers for Norway spruce that would be used for three further purposes: (1) assembly improvement and assessment, via the estimation of a scaffold-anchored genetic map and the inclusion of probe pairs stradling contig joins within a scaffold, to test scaffolding decisions made during assembly (Sahlin et al., 2014); (2) trait-based association studies, to understand the architecture of quantitative traits and to assist the design of artificial selection experiments for breeding; and (3) population genomic studies, to understand the evolutionary forces that have shaped genome structure and variation. We chose to develop two partially-overlapping sets of sequence probes assayed against different sets of tissue samples. For purpose (1), we developed probes against haploid megagametophyte tissues related to the sequenced tree Z4006, which limits the general usefulness of the marker set but improves its utility for the assembly. For purposes (2) and (3), we developed probes against diploid needle samples from throughout the range of Norway spruce.

Considering the high repetitive content within the Norway spruce genome, including high conservation in some repetitive element families (Zuccolo et al. 2015), we focused probe development on exons within genes, as delineated via the annotation accompanying *P. abies* assembly version 1.0. The ~67,000 annotated nuclear gene models (*ab initio*-predicted protein-coding loci) included in the assembly are divided into three categories designating the relative degree of support for the gene model, based on alignment of supporting evidence provided by non-*P. abies* protein or EST sequences: high-confidence (HC) gene models (39.7%), which were covered >70% of the model length; medium-confidence (MC) gene models (48.1%), covered 30-70%; and low-confidence (LC) gene modules (12.2%), covered <30%. See Nystedt et al. (2013) for further details of gene model development and confidence categories. During probe development, we favoured exons of the HC gene models but also included subsets of MC and LC gene models. The goal was to place a probe within each HC gene model, and where practical two probes/HC gene, resulting in approximately 40,000 probes. We did not design the probe sets to cover complete exons nor did we design probes against all exons of each gene.

In this study, we discuss the development and evaluation of sequence sequence capture probe sets developed against haploid and diploid tissues in Norway spruce.

## Methods

Figure 1 provides an overview of the probe design workflow.

**Figure 1:**
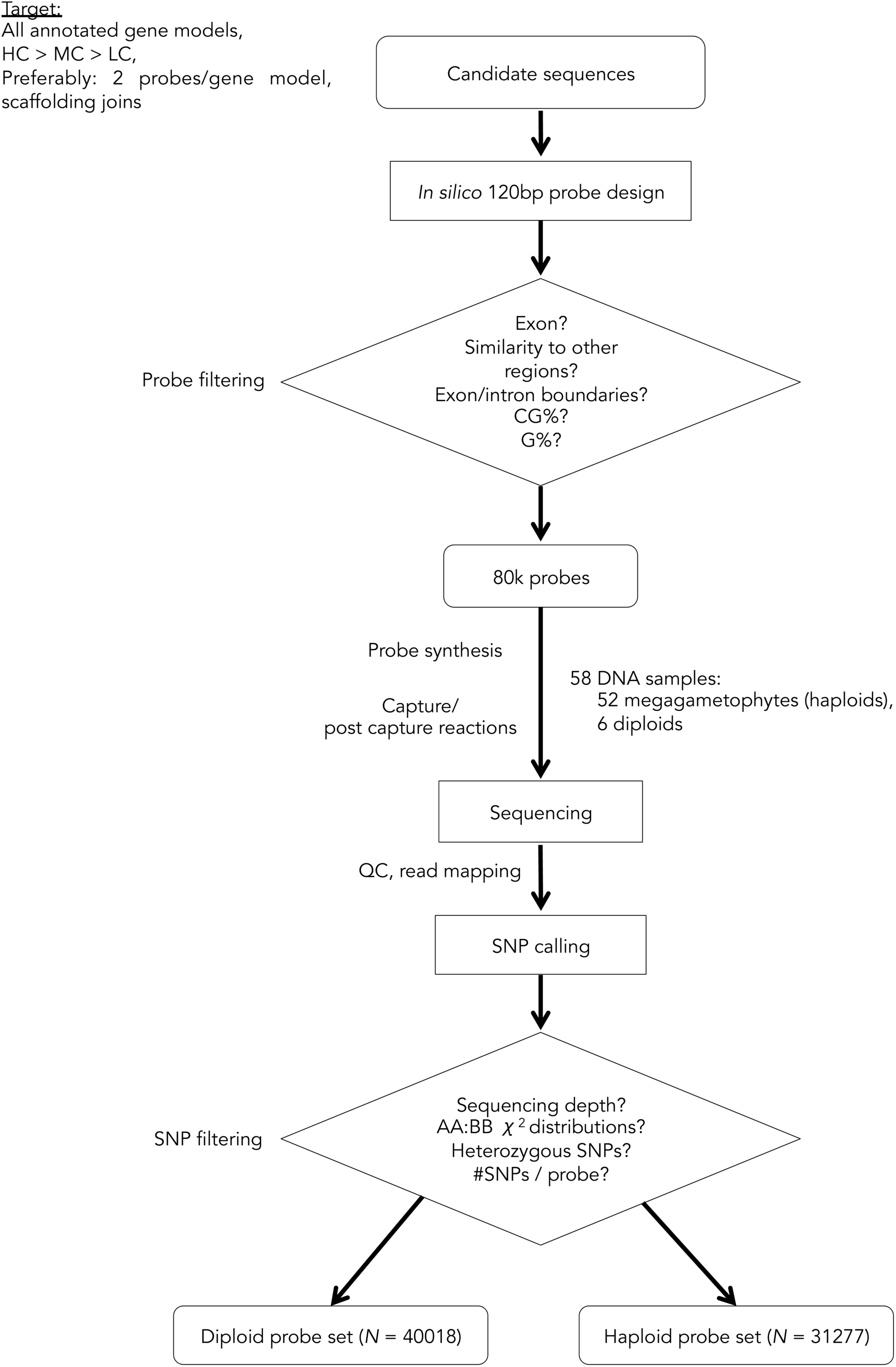
Probe selection workflow.

### Probe design on candidate sequences

Candidate sequences for probe design were created based on RNA-seq and scaffolding decisions of Pabies 1.0. Our goal was to design probes on either side of scaffolding joints, in order to assist the genome assembly. To achieve that, the sequences corresponding to HC, MC and LC gene models were selected for probe design, with priority given subsequently to HC and MC categories. All possible probes (120nt) were designed in silico on the candidate genes, with start-end coordinates provided as target subsequences. We included separate subsequences because these represented separate contigs prior to RNA-seq scaffolding or paired-end/mate-pair scaffolding in *P.abies* 1.0. Ideally, we aimed at having separate probes targeting each subsequence to test whether the RNA-seq and paired-end/mate-pair scaffolding decisions were made correctly. All RNA-seq scaffolding events that involved annotated genes were included in the candidate subsequences. As the number of paired-end/mate-pair scaffolding events that involved annotated genes was considerably larger, a random subsample of 15% of these events was included in subsequence delineation. From the total of all possible probes within the candidate sequences, filters were applied to select a set of 80,000 probes that were used for hybridization in the pilot experiment. First, sequencing-level removed probes with extreme GC content (<0.2 and >0.6), high G content (>0.2) and with long homopolymers (>7). Next, probes falling on exon-exon boundaries (as indicated by information from available genome annotation) were removed. Finally, probes were mapped to the genome and chloroplast sequences. Probes mapping to the chloroplast and those aligning to more than one position (90% identity for 90% or the length) were excluded. From the resulting probes, a maximum of two probes per subsequence were chosen to comprise the final probe set.

### Plant material and DNA extraction

Haploid genomic DNA was extracted from 52 megagametophytes. The megagametophytes were excised from open pollinated seeds of Z4006 ramets (Z4006: the Norway spruce reference sequence individual), under the microscope in order to avoid diploid tissue. DNA was extracted with the NucleoSpin^®^ Plant II kit, (Macherey-Nagel, http://www.mn-net.com). After several modifications of the manufacture’s recommended protocol, we achieved the highest concentration of DNA from megagametophytes (mean concentration of 40.6 ng/µl) by grinding the megagametophytes together with the extraction buffer in an electric grinder. Diploid genomic DNA was extracted from lyophilized leaves of six individuals that span a large range of the geographic distribution of *Picea abies*. The six individuals were sampled in Russia, Poland, Belarus, Romania and Southern Sweden, including the reference genome sequenced individual Z4006.

### Library preparation and Target enrichment

Extracted DNA was submitted for RAPiD Genomics (USA) where DNA library preparation and capture sequencing were performed. The concentration of the extracted DNA was estimated with PicoGreen dsDNA quantification assay (ThermoFisher Scientific, USA) and DNA integrity was analyzed by visualizing the DNA on a 0.8% w/v agarose electrophoresis gel. Libraries compatible with Illumina sequencing were prepared with varying starting amounts of DNA, depending on the yield of the DNA extraction, between 450-500 ng. The DNA was mechanically sheared to a mean fragment size of 300bp, followed by repair of the ends of the molecules, phosphorylation and adenylation. Illumina TruSeq equivalent adapters suited for sequencing were ligated on each side of the molecules containing different 8bp indexes (i7). The libraries were amplified with 14 cycles of PCR and the resulting libraries were quantified with PicoGreen. The set of 80,000 probes synthesized as 120 nt RNA molecules were hybridized to a pool containing a total of 500 ng from 8 equimolarly combined libraries following Agilent’s SureSelect Target Enrichment System (Agilent Technologies). The enriched libraries were sequenced on one lane of Illumina HiSeq 2000 and two lanes of HiSeq 500 high-output instruments on a 1x100bp and 1x75bp sequencing mode, respectively.

### Probe evaluation

Reads from sequences captured with the 80,000 pilot probes were mapped to the *P. abies* 1.0 probe-containing scaffolds and variants at each probe site were called with FreeBayes (Garrison and Marth 2012) followed by filtering for heterozygosity and expected 1:1 segregation ratio. Regions ±300 bp around each probe site were included to capture more variants. Further evaluation and filtering was applied to this variant set to select the initial megagametophyte and diploid probe sets.

Probe context was evaluated by comparison with the *P. abies* 1.0 genome annotation. For each probe set, we used BEDTools (Quinlan and Hall 2010) to intersect 120-bp probe sites with gene models. We identified four separate features: (1) exonic sequence, marked as CDS within the genome annotation; (2) intronic sequence, between separate CDS sequences; (3) UTR-like sequences, which were not annotated directly within the genome annotation but which we inferred as being 1-500 bp upstream of the annotated translation Start site or 1-500 bp downstream of the annotated translation Stop site; (4) exon-intron splice sites.

The selected probe sets were used for additional megagametophyte and diploid sequencing. After sequence delivery, probe sites were subject to further evaluation following additional read-mapping with BWA, duplicate marking with Picard, 1.127, and GATK 3.4.0 for realignment and variant-calling with both UnifiedGenotyper and HaplotypeCaller, with a focus on developing reliable variant sites. Read depth, duplicate and multiply-mapped reads, unusual coverage depth, variant quality and breadth of probe site coverage were examined programmatically. For a collection of probe sites, the results of multiple read-mapping and variant-calling options were subject to direct examination in IGV.

## Results and Discussion

### Probe design (pre-sequencing)

Prior to probe design, 66,632 gene models in the P. abies 1.0 genome annotation were examined and 76,144 candidate sequences were selected for potential probe design, with mean 1.14 candidates/gene model and a total length of 166 Mbp (Table 1A). Within these candidate sequences, 403,357 potential 120-bp probe sites were identified (Table1B). After initial screening and redundant probe removal (≤ 2 probes/sequence) these were reduced to 80,000 total pilot probes, with 34,761 gene models and 32,495 separate scaffolds containing at least one pilot probe site (Table 1B).

**Table 1:**
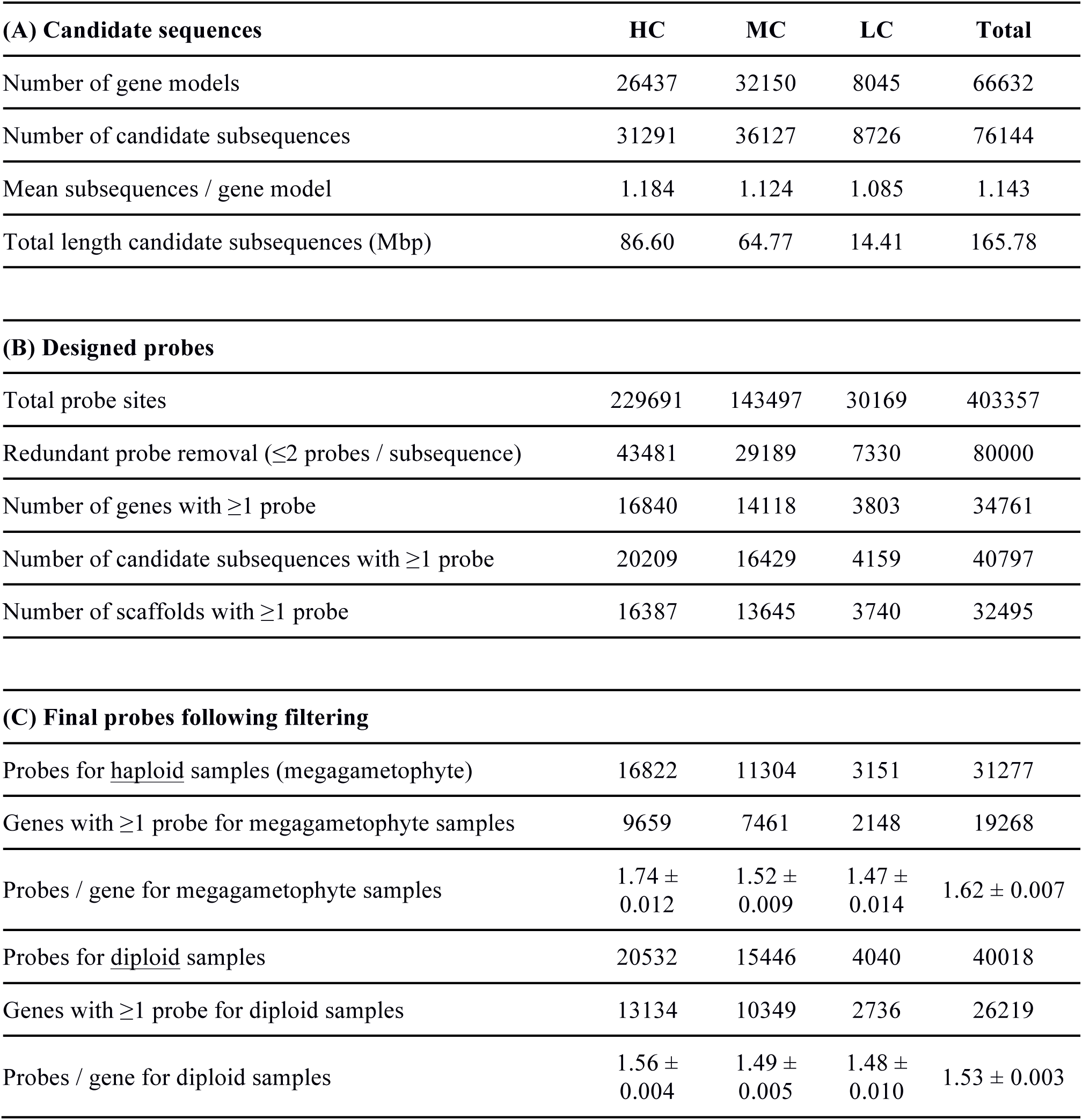
Basic probe targeting statistics vs. Norway spruce 1.0 genome sequence. HC, MC, LC = High-, Medium- and Low-Confidence gene models, respectively; see text for details.

| **(A) Candidate sequences** | **HC** | **MC** | **LC** | **Total** |
| --- | --- | --- | --- | --- |
| Number of gene models | 26437 | 32150 | 8045 | 66632 |
| Number of candidate subsequences | 31291 | 36127 | 8726 | 76144 |
| Mean subsequences / gene model | 1.184 | 1.124 | 1.085 | 1.143 |
| Total length candidate subsequences (Mbp) | 86.60 | 64.77 | 14.41 | 165.78 |
| **(B) Designed probes** |  |  |  |  |
| Total probe sites | 229691 | 143497 | 30169 | 403357 |
| Redundant probe removal ($\leq 2$ probes / subsequence) | 43481 | 29189 | 7330 | 80000 |
| Number of genes with $\geq 1$ probe | 16840 | 14118 | 3803 | 34761 |
| Number of candidate subsequences with $\geq 1$ probe | 20209 | 16429 | 4159 | 40797 |
| Number of scaffolds with $\geq 1$ probe | 16387 | 13645 | 3740 | 32495 |
| **(C) Final probes following filtering** |  |  |  |  |
| Probes for <u>haploid</u> samples (megagametophyte) | 16822 | 11304 | 3151 | 31277 |
| Genes with $\geq 1$ probe for megagametophyte samples | 9659 | 7461 | 2148 | 19268 |
| Probes / gene for megagametophyte samples | $1.74 \pm 0.012$ | $1.52 \pm 0.009$ | $1.47 \pm 0.014$ | $1.62 \pm 0.007$ |
| Probes for <u>diploid</u> samples | 20532 | 15446 | 4040 | 40018 |
| Genes with $\geq 1$ probe for diploid samples | 13134 | 10349 | 2736 | 26219 |
| Probes / gene for diploid samples | $1.56 \pm 0.004$ | $1.49 \pm 0.005$ | $1.48 \pm 0.010$ | $1.53 \pm 0.003$ |

### Probe evaluation against sequenced samples

An initial set of sequence capture results using both haploid megagametophytes and diploid needle tissue was produced with these 80,000 pilot probes, and variants were called within the probe site using FreeBayes (Garrison and Marth 2012). After initially finding low numbers of heterozygous variants that segregated at ~1:1 ratio, queried sites were expanded ±300 bp of the boundaries of each probe site. This recovered sufficient variants to proceed with selection among pilot probes.

After evaluation of probe site variant qualities in haploid and diploid read sets separately, two partially overlapping sets of final probes was selected for further sequence capture: 31,277 sites for haploid megagametophytes and 40,018 sites for diploid needle tissues (Table 1C). The initial set of 40,000 diploid probe sites was expanded to 40,018 by adding 18 sites covering some genes of interest that were filtered out in earlier screening. In the megagametophyte set, which will be used primarily for construction of genetic maps for assembly evaluation and improvement, 19,268 gene models were included, with an average of 1.62 ± 0.007 probes/model (Table 1C). The diploid probe set covered 26,219 gene models with an average of 1.53 ± 0.003 probes/model (Table 1C).

### Probe context

For each of the final probe sets, we evaluated the context of probe sites vs. gene models in the *P. abies* 1.0 genome annotations. The probes were designed to be used against DNA resulting from whole-genome extractions, so exonic, intronic and possible UTR sequences are all possible within probe sites, as are exon-intron splice sites. Within the 120-bp probe sites of the 31,277 megagametophyte probes, a total of 1600.4 Kbp of exonic sequence was covered, 2152.9 Kbp of intronic sequence was covered, 28.5 Kbp of UTR-like sequence was covered, and 6195 exon-intron boundaries were covered (Table 2). For the 40,018 diploid probes, a total of 2331.1 Kbp of exonic sequence was covered, 2470.9 Kbp of intronic sequence was covered, 40.7 Kbp of UTR-like sequence was covered, and 9119 exon-intron boundaries were covered (Table 2).

**Table 2:** Probe context vs. Norway spruce 1.0 genome sequence. HC, MC, LC = High-, Medium- and Low-Confidence gene models, respectively; see text for details.

| **PROBE CONTEXT: 31K megagametophyte** | **HC** | **MC** | **LC** | **Total** |
| --- | --- | --- | --- | --- |
| Exonic sequence (Kbp) | 873.7 | 547.2 | 179.6 | 1600.4 |
| Intronic sequence (Kbp) | 1145.0 | 809.3 | 198.6 | 2152.9 |
| UTR-like sequence (Kbp) (outside exons/introns, $\pm 500$bp of Start/Stop) | 11.5 | 14.5 | 2.6 | 28.5 |
| Exon-Intron splice sites under probes | 3142 | 2416 | 637 | 6195 |
| **PROBE CONTEXT: 40K diploid** |  |  |  |  |
| Exonic sequence (Kbp) | 1298.9 | 810.2 | 222.0 | 2331.1 |
| Intronic sequence (Kbp) | 1164.8 | 1043.3 | 262.8 | 2470.9 |
| UTR-like sequence (Kbp) (outside exons/introns, $\pm 500$bp of Start/Stop) | 17.4 | 20.0 | 3.3 | 40.7 |
| Exon-Intron splice sites under probes | 4371 | 3810 | 983 | 9119 |

### Evaluation of variants within probe sites and switch to whole-genome mapping

To further evaluate the selected probe sets, sequence capture reads from 58 megagametophyte samples and 6 diploid samples were aligned to probe-containing scaffolds using BWA-MEM (Li 2013), followed by duplicate marking with Picard and realignment with GATK and variant calling with both UnifiedGenotyper and HaplotypeCaller in GATK. and variants were called following indel realignment. Following further analysis of the two probe sets, the numbers of filtered variants still seemed unusually low. Direct examinations of read mappings and variant calls within selected probe sites using IGV indicates that at some probe sites, the mapped reads included reads clearly from outside the probe site, as indicated by lower mapping quality, differences that did not clearly belong to one or two haplotypes in the megagametophyte or diploid probe sets, and disagreements among methods in variant presence and quality.

Considering these observations together, we hypothesised that these problems were caused by occasional promiscuous capture of sequences from outside probe sites and more importantly, from genome sequences not included in the set of probe-containing scaffolds. The correct alignment target of such external sequences would not be present in the probe-containing scaffolds, so instead the reads would be mapped to the best available sites. This is likely to be encountered any time the potential source of reads exceeds the reference to which they are being aligned.

To overcome this problem, we switched to aligning sequence capture reads to the complete *P. abies* 1.0 genome assembly. This assembly lacks ~7.5 Gbp from the estimated 19.6 Gbp in the complete genome, but much of the missing sequence is likely to to be repetitive (Nystedt et al. 2013) and thus excluded by our probe design. After mapping to the complete assembly, we then restricted the read alignments to just those that were found on probe-containing scaffolds. This resulted in the loss of ~5% of sequenced reads for each sample (Figure 2).

**Figure 2:**
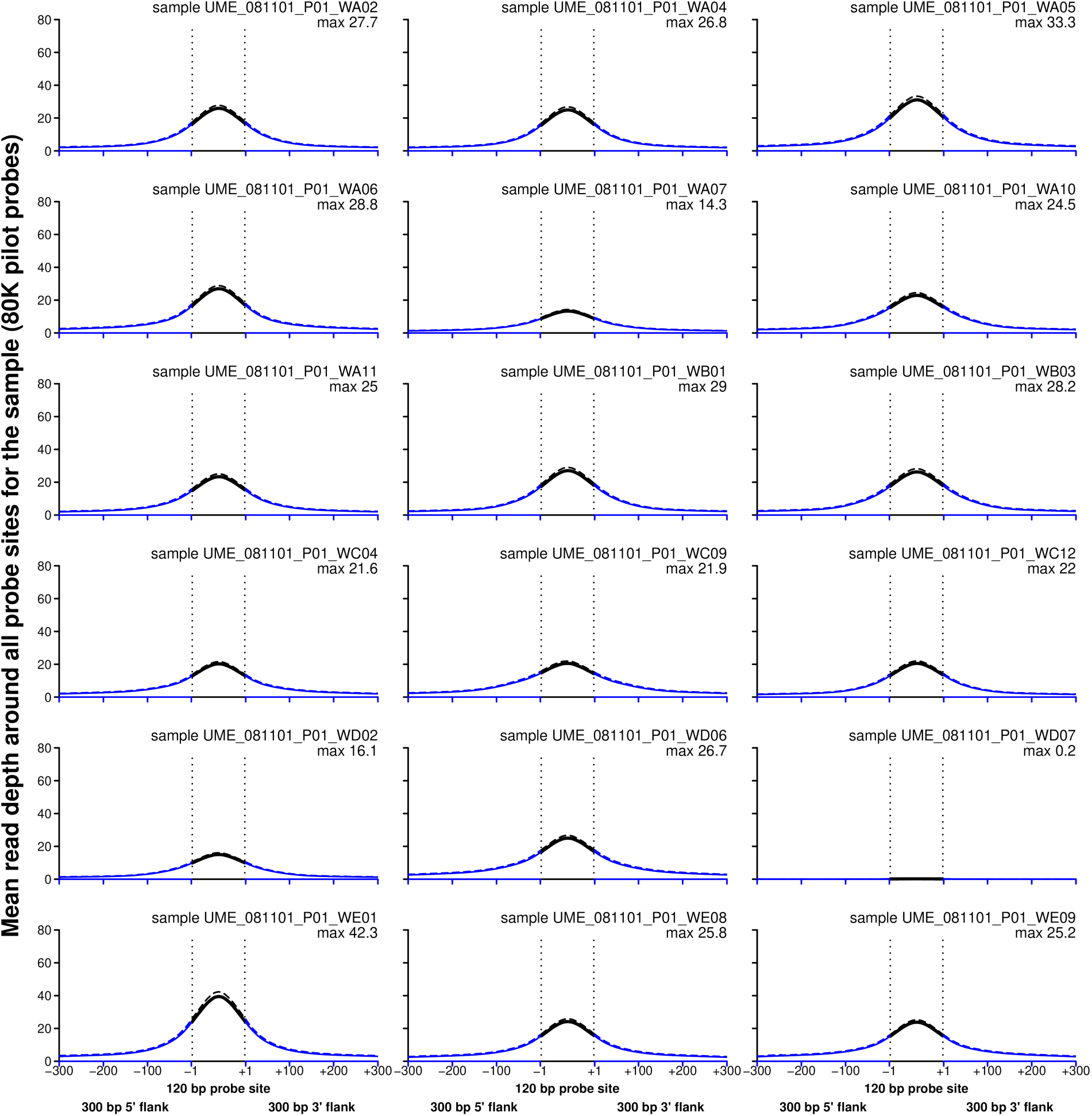
Mean read depth over all 80,000 pilot probe sites for each of a selection of 18 pilot samples. Each panel shows mean read depth from libraries derived from sequence capture across all probe sites (y-axis) within a window around all probe sites (x-axis). The 120-bp probe site is bounded by vertical dotted segments and read depth within the probe site proper is shown in black lines. Also included is mean read coverage 300 bp up- and downstream of the probe site (blue lines). Mean read depth is shown for two methods of read alignment: solid lines show depth when reads are mapped to the complete *Picea abies* 1.0 genome; and dashed lines show depth when reads are mapped only to the probe-containing scaffolds. Also shown is the sample name and maximum mean read depth within the 720-bp window shown.

At some probe sites, the difference was quite dramatic (Figure 3). Most of the probe sites show relatively little difference in read coverage when reads are mapped to the whole genome or to the restricted set of probe-containing scaffolds. At some illustrated probes, for example probes 33254, 42123, 44589, 50875 and 58730, 25% or more of read coverage within and around the probe site was mapped elsewhere when mapped against the full 1.0 genome assembly (Figure 3).

**Figure 3:**
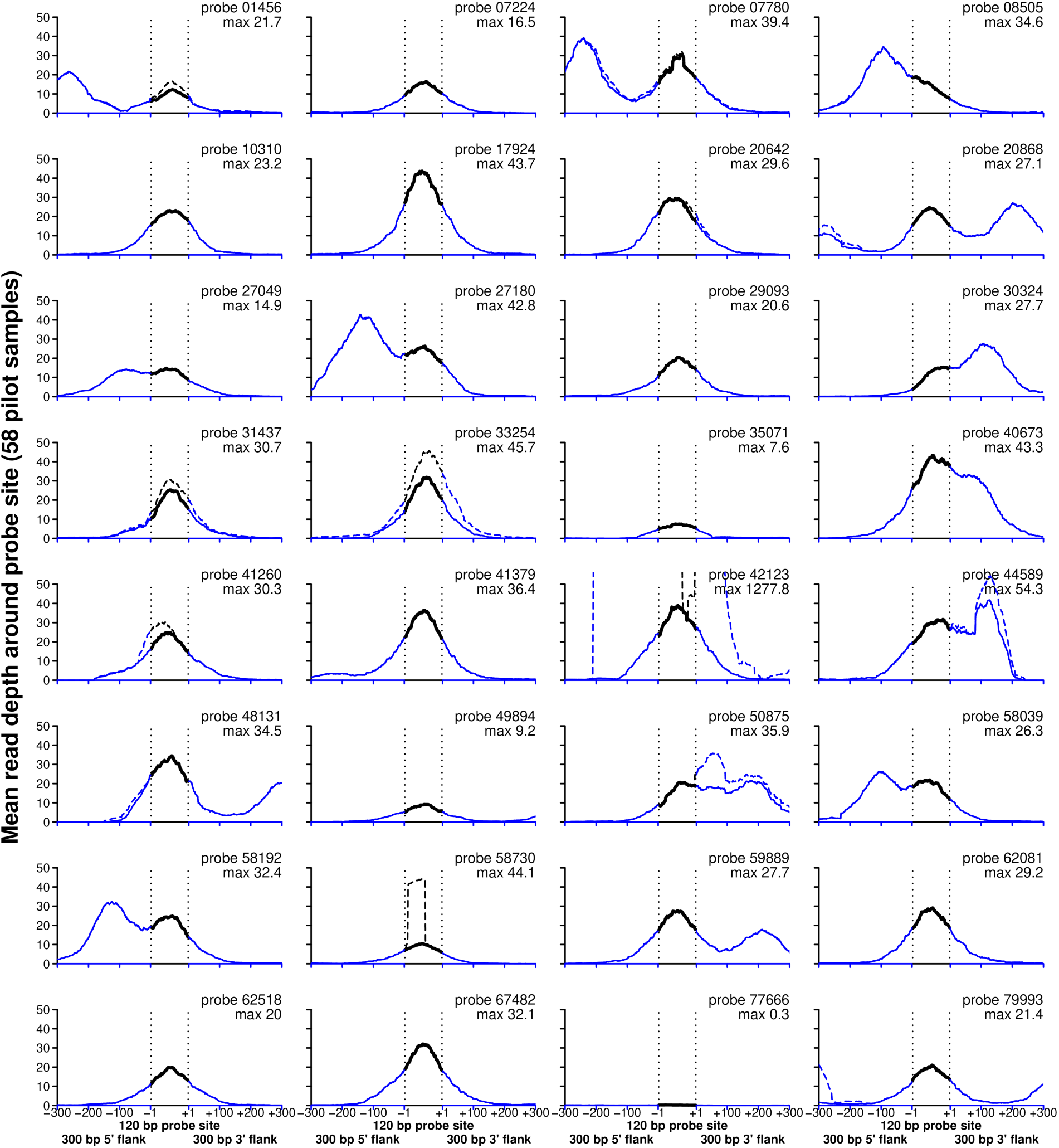
Mean read coverage over each of a selection of 32 probe sites across all 58 pilot samples. Each panel shows mean read depth from libraries derived from sequence capture across all 58 pilot samples (y-axis) within a 720-bp window around a single focal probe site (x-axis). Otherwise the colouring and plotting is as described for Figure 2. As for Figure 2, mean read depth is shown for two methods of read alignment: solid lines show depth when reads are mapped to the complete *Picea abies* 1.0 genome; and dashed lines show depth when reads are mapped only to the probe-containing scaffolds. Coverage for the latter, which offers limited control against off-target sequence capture, may exceed the depth-50 limit of the y-axis, as for probe 42123. Also shown is the pilot probe designation and maximum mean read depth across samples within the 720-bp window shown for that probe.

### Probe precision

The average read coverage across all probe sites within individual samples quite clearly centred on the probe site proper, with symmetric drop-off in coverage on either site of the 120-bp site (Figure 2). A closer look at a selection of individual probe sites across samples reveals that differences among probe sites proper can be quite dramatic (Figure 3). Most probe sites reveal good targetting of the site, and most sequences mapped within a larger 720-bp window which includes 300 bp up- and downstream of the probe site are from the probe site or within 100 bp of the probe site. In light of these results, we decided to accept variants found within the probe site or within 100 bp up- or downstream of the probe site when selecting variants called from large-scale megagametophyte and diploid sequence capture.

Several probe sites show a bimodal read depth within the 720-bp window (Figure 3). It is not immediately clear why this would be the case, but it is not rare and it is consistent across samples. In some cases extra-site reads are from other sites in the genome (e.g., probes 42123) but in most, the coverage persists. This may represent some local bias during DNA fragmentation or variation in probe capture kinetics.

## Availability of probe sequences

The three sets of probe sets described here – the pilot set, the megagametophyte set, and the diploid set – are available at https://github.com/douglasgscofield/pubs/tree/master/Vidalis-et-al-1.

## Acknowledgements

We are grateful to the captain and the crew of LH2414 for bringing AV safely to Sweden in January 2016. AV was supported in part by the Stiftelsen Gunnar och Birgitta Nordins fond through the Kungl. Skogs-och Lantbruksakademien. The Norway spruce genome project was supported by the Knut and Alice Wallenberg Foundation. The computations were performed on resources provided by SciLifeLab and SNIC at the Uppsala Multidisciplinary Center for Advanced Computational Science (UPPMAX) under project b2010042.

## References

Clark MJ, Chen R, Lam HYK, Karczewski KJ, Chen R, Euskirchen G, Butte AJ, Snyder M. 2011. Performance comparison of exome DNA sequencing technologies. Nat Biotech 29:908–914.

Davey JW, Blaxter ML. 2010. RADSeq: next-generation population genetics. Briefings in Functional Genomics 9: 416–423.

Fu W, O’Connor TD, Jun G, Kang HM, Abecasis G, Leal SM, Gabriel S, Altshuler D, Shendure J, Nickerson DA, et al. 2013. Analysis of 6,515 exomes reveals the recent origin of most human protein-coding variants. Nature 493: 216–220.

Garrison E, Marth G. 2012. Haplotype-based variant detection from short-read sequencing. arXiv:1207.3907 [q-bio.GN].

Li H. 2013. Aligning sequence reads, clone sequences and assembly contigs with BWA-MEM. arXiv:1303.3997v1 [q-bio.GN].

Neale DB, Wegrzyn JL, Stevens KA, Zimin AV, Puiu D, Crepeau MW, Cardeno C, Koriabine M, Holtz-Morris AE, Liechty JD, et al. 2014 Decoding the massive genome of loblolly pine using haploid DNA and novel assembly strategies. Genome Biol 15: R59.

Neves LG, Davis JM, Barbazuk WB, Kirst M. 2013. Whole-exome targeted sequencing of the uncharacterized pine genome. The Plant Journal 75: 146–156.

Nystedt B, Street NR, Wetterbom A, Zuccolo A, Lin Y-C, Scofield DG, Vezzi V, Delhomme N, Giacomello S, Alexeyenko A, et al. 2013. The Norway spruce genome sequence and conifer genome evolution. Nature 497: 579–584.

Quinlan AR, Hall IM. 2010. BEDTools: a flexible suite of utilities for comparing genomic features. Bioinformatics 26: 841–842.

Sahlin K, Vezzi F, Nystedt B, Lundeberg J, Arvestad L. 2014. BESST: Efficient scaffolding of large fragmented assemblies. BMC Bioinformatics 15:281.

Shigemizu D, Momozawa Y, Abe T, Morizono T, Boroevich KA, Takata S, Ashikawa K, Kubo M, Tsunoda T. 2015. Performance comparison of four commercial human whole-exome capture platforms. Scientific Reports 5: 12742.

Suren H, Hodgins KA, Yeaman S, Nurkowski KA, Smets P, Rieseberg LH, Aitken SN, Holliday JA. 2016. Exome capture from the spruce and pine giga-genomes. Molecular Ecology Resources 16: 1136–1146.

Wang J, Street NR, Scofield DG, Ingvarsson PK. 2016. Variation in linked selection and recombination drive genomic divergence during allopatric speciation of European and American aspens. Molecular Biology and Evolution 33: 1754–1767.

Wang J, Ding J, Tan B, Robinson KM, Michelson IH, Johansson A, Nystedt B, Scofield DG, Nilsson O, Jansson S, Street NR, Ingvarsson PK. 2017. Natural variation at FLOWERING LOCUS T2 mediates local adaptation in a key life history trait in European aspen. bioRxiv 178921. doi: https://doi.org/10.1101/178921

Zhou Y, Tao S, Chen H, Huang L, Zhu X, Li Y, Wang Z, Lin H, Hao F, Yang Z, Wang L and Zhu X. 2014. Exome sequencing analysis identifies compound heterozygous mutation in ABCA4 in a Chinese family with Stargardt disease. PLoS ONE 9: e91962.

Zuccolo A, Scofield DG, De Paoli E, Morgante M. 2015. The Ty1-*copia* LTR retroelement family PARTC is highly conserved in conifers over 200 MY of evolution. Gene 568: 89–99.

